# Grant reviewer perceptions of the quality and effectiveness of panel discussion

**DOI:** 10.1101/586685

**Authors:** Stephen A. Gallo, Karen B. Schmaling, Lisa A. Thompson, Scott R. Glisson

**Affiliations:** Scientific Peer Advisory and Review Services, American Institute of Biological Sciences, Herndon, VA; Washington State University, Vancouver, WA

**Keywords:** Peer Review, Discussion, Research Funding, Grant Applications, Survey

## Abstract

**Background:** Funding agencies have long used panel discussion in the peer review of research grant proposals as a way to utilize a set of expertise and perspectives in making funding decisions. Little research has examined the quality of panel discussions and how effectively they are facilitated.

**Methods:** Here we present a mixed-method analysis of data from a survey of reviewers focused on their perceptions of the quality and facilitation of panel discussion from their last peer review experience.

**Results:** Reviewers indicated that panel discussions were viewed favorably in terms of participation, clarifying differing opinions, informing unassigned reviewers, and chair facilitation. However, some reviewers mentioned issues with panel discussions, including an uneven focus, limited participation from unassigned reviewers, and short discussion times. Most reviewers felt the discussions affected the review outcome, helped in choosing the best science, and were generally fair and balanced. However, those who felt the discussion did not affect the outcome were also more likely to evaluate panel communication negatively, and several reviewers mentioned potential sources of bias related to the discussion. While respondents strongly acknowledged the importance of the chair in ensuring appropriate facilitation of the discussion to influence scoring and to limit the influence of potential sources of bias from the discussion on scoring, nearly a third of respondents did not find the chair of their most recent panel to have performed these roles effectively.

**Conclusions:** It is likely that improving chair training in the management of discussion as well as creating review procedures that are informed by the science of leadership and team communication would improve review processes and proposal review reliability.

## Background

The US National Institutes of Health (NIH), like many major research funders, utilizes a “long standing and time-tested system of peer review to identify the most promising biomedical research (Liaw 2017; NIH 2018).” However, many reports of poor inter-rater reliability suggest a high degree of subjectivity to the process (Cole, 1981; Chichetti, 1991; Fogelholm 2012; Pier et al., 2018). One common procedure used to mitigate this subjectivity is to discuss each proposal at a meeting of the entire review panel, utilizing a larger set of expertise and perspective than just that of the 2-3 reviewers that are explicitly assigned to read and evaluate the proposal pre-meeting (Brown 1993, Pier 2017). In this system, final scores are typically generated from the average post-discussion scores of all assigned and unassigned reviewers (without conflicts of interest), in an attempt to ensure all available expertise is brought to bear on the final evaluation of the proposal (NIH 2018).

Despite this intended goal, several studies have reported that discussion can have a somewhat limited effect on the final scoring of proposals (Obrecht, 2007, Martin 2012, Carpenter 2015, Fogelholm 2012, Pier 2017), with some studies estimating that a proposal’s funding status (score above or below the funding line) is shifted from pre-to post-discussion for only 10-13% of proposals (Fogelholm 2012, Carpenter 2015). While some studies (Fleurence et al., 2014) suggest that most discussions do yield changes in assigned reviewers’ scores (which in general move closer together, narrowing the range of scores), the magnitude of scoring shifts after discussion are typically relatively small and are even smaller (with shorter discussion times) in teleconference panels compared to face-to-face panels (Carpenter 2015, Pier 2015). It is known that face-to-face communication is a richer channel than other virtual alternatives (Cooke et al., 2015), and it is likely that the quality of panel communication plays an important role in how influential the panel discussions are on scoring.

Interestingly, it has been observed that scoring shifts are correlated with instances of panel discourse “in which reviewers made explicit references to the scoring habits of fellow panelists,” which have been referred to as score calibration talk (Pier 2017). These scoring shifts are particularly related to discussion where reviewers were being held accountable by other reviewers for how they calibrated their score relative to the descriptors in their written critiques. For instance, “your comments are meaner than your score” (Pier 2017). However, it is not clear how well these types of interactions are promoted by the chair or if all reviewers (especially unassigned reviewers) feel they have the opportunity to present such opinions. In fact, others have noted that, although the goal of convening panels is to bring a range of expertise to bear on the evaluation of a research proposal, opportunities are limited for dialogues between reviewers of different expertise (Langfeldt, 2001).

In addition, no studies have examined the effectiveness of panel discussion in highlighting where differences in opinion occur, so that unassigned reviewers are well-informed in their final scoring decisions. While, in general, unassigned reviewers’ scores closely mirror assigned reviewers’ scores (Martin 2010), some assigned reviewers exert more influence than others, which may be due to “expertise, authority and debating, persuasion, or argumentation skills” (Fogelholm 2012). The chair’s role is to facilitate discussion, moderate individual personalities, and provide a fair and balanced presentation of each proposal under evaluation to unassigned reviewers. However, it is not clear if all chairs effectively fulfill these responsibilities, and studies have not been undertaken to evaluate the effectiveness of panel chairs in stewarding peer review panel proceedings. Such evaluations are, therefore, needed to better understand the effectiveness of panel chairs, particularly since this type of information cannot be derived from peer review scores.

Little data are available regarding reviewers’ perceptions of panel discussions. Some favorable reviewer perceptions have, however, been reported by NIH, specifically that 81% agreed or strongly agreed that “scientific discussions supported the ability of the panel to evaluate the applications being reviewed” (NIH CSR 2015), but the general nature of this statement does not provide a finer-grained understanding of panel discussions. Details of interest include the level of participation in discussion, whether discussion helps to clarify opinions, how well the chair facilitates this discussion, and how well unassigned reviewers can make informed decisions based on the discussion. Moreover, it would be pertinent to determine if reviewers feel that the discussions affect the outcome of proposal reviews.

To address these gaps, we developed a survey focused on reviewer perceptions of their most recent panel meeting experience and distributed it to a diverse group of research scientists. Two publications have resulted from analyses of the survey responses (Gallo et al., 2018, Gallo et al., 2019), but neither publication focused on the section of the survey that addressed perceptions of the quality and facilitation of panel discussion and their impact on review outcomes. To examine these topics, feedback from the surveyed scientists were summarized regarding the quality and facilitation of their most recent panel discussions with the goal of developing a better understanding of reviewer perceptions of panel effectiveness to help inform the future implementation of review formats and procedures.

## Methods

### Survey

This study involved a diverse group of biomedical research scientists who responded to a survey. The survey was reviewed by the Washington State University Office of Research Assurances (Assurance# FWA00002946) and granted an exemption from IRB review consistent with 45 CFR 46.101(b)(2). Participants were free to choose whether or not to participate in the survey and consented by their participation. They were fully informed at the beginning of the survey as to the purpose for this research, how we acquired their email address, and the importance and intended use of the data. The general survey methodology has been described in two other manuscripts (Gallo et al., 2018; Gallo et al., 2019). The original survey contained 60 questions and was divided into 5 subsections; data from only 3 sections are presented in this manuscript to address the issue of peer review panel discussion quality: 1. grant submission and peer review experience; 2. reviewer attitudes toward grant review; and 3. peer review panel meeting proceedings. The questions regarding discussion quality included here were not analyzed in the previous publications, although other aspects, such as review frequency and reviewer preference, were reported previously.

The questions examined had either nominal (Yes/No) or ordinal (Likert rating) response choices. For example, on a scale of 1-5 (1=most definitely, 5=not at all), did the grant application discussions promote the best science? However, respondents were also given the choice to select no answer/prefer not to answer. At the end of each section, respondents could clarify their answers in a free form text box. A full copy of the peer review survey is available in the **S1 File in the Supporting Information**. The raw, anonymized data are also available (https://doi.org/10.6084/m9.figshare.8132453.v1).

As described in previous publications, the survey was sent out in September of 2016 to 13,091 individual scientists from the American Institute of Biological Science’s (AIBS’s) database through the use of Limesurvey (© Hamburg, Germany), which de-identified the responses from respondents. AIBS’s database has been developed over several years to help AIBS recruit potential reviewers for evaluation of biomedical research applications for a variety for funding agencies, research institutes and non-profit research funders. Most of these reviews are non-recurring and scientists are recruited based on matching expertise to the topic areas of the applications. The individuals invited to this survey were either reviewers for AIBS (26%) or had submitted an application as a PI that was reviewed by AIBS (62%) or both (12%). Depending on the question, respondents were asked to focus on either the most recent peer review or reviews that occurred in the last 3 years.

### Procedure and Data Summarization

The survey was open for two months; responses were then exported and analyzed using Stat Plus software. For this paper, participants were included only if they completed the survey and included an answer for questions 2e and 2f, which focused on whether they had participated in a peer review panel in the last three years and, if so, how often. Thus, all questions included in this analysis were focused on reviewer experiences. Reviewers were asked questions related to the qualities of panel discussion. Mean and percentage comparisons were analyzed using non-parametric tests (e.g. Mann-Whitney, chi-square tests), due to the highly skewed ordinal distributions (most are >1.0). Standard 95% confidence intervals (CI) were calculated for the Likert responses (for proportion data, binomial proportion confidence intervals were calculated). Effect size (d) was calculated via standardized mean difference for all comparisons. Differences between groups were considered significant if there was either no overlap in CI or if there was overlap yet a test for difference indicated a significant result (p<0.01).

All comments made by respondents from the survey related to the Peer Review Panel Meeting Proceedings section were extracted. All quotes were then grouped according to which of the six specific questions analyzed in this manuscript they most closely addressed. Multiple groupings per quote were permissible. Many of the quotes in this section were not relevant to the six questions or the survey section (e.g. their level of participation) and were therefore not considered for inclusion in this analysis. The remaining quotes were then examined for common themes, and when multiple quotes related to a similar idea were identified, quotes were selected for inclusion in the manuscript that were well-written, clear and specific relative to the survey question. If contrasting views on the same theme were expressed, care was taken to ensure that both quotes were included in the manuscript. If only part of a respondent’s comment was relevant to one of the six questions, only the relevant portion was included.

## Results

### Response Rate, Demographics and Review Setting

Of the 13,091 individuals contacted for this survey, 1231 responded, giving a 9.4% response rate. Of the 1231 respondents, only 874 of these completed questions 2e and 2f, 671 (77%) of whom indicated they had recently reviewed on a panel in the last 3 years. These 671 reviewer respondents formed the core group upon which the current analyses are based. Of these, free text answers were provided by 159 respondents. A total of 29 quotes were used in relation to specific survey questions; these are presented below.

Demographics were analyzed in detail in previous publications: respondents were 66% male, 80% PhD, and 69% in a late career stage (e.g. tenured full and emeritus professorship) with the majority being age 50 or older (75%; median age 55), Caucasian (76%) and working in academia (81%). These respondents participated in an average of 4.0 ± 0.08 panel meetings over the last 3 years.

### Discussion Quality

Utilizing the expertise of the whole panel is one of the rationales for discussing proposals at the meetings. As one respondent put it:

> “Discussions can occasionally develop a herd mentality, so having multiple perspectives represented and facilitation to hear all voices is crucial.”

The vast majority (92% [90%-95%]) of reviewers felt the panel discussions involved reviewer participation, although participation may be in reference to the engagement of assigned reviewers. Some respondents specifically mentioned participation from unassigned reviewers, although their participation in discussion was not always at a high frequency:

> “Overall, discussion was driven by the assigned reviewers. Sometimes an interested non-assigned reviewer would get involved. Sometimes, non-assigned reviewers would ask clarifying questions.”
>
> “Non-assigned reviewers rarely ask questions or comment.”

70% of all reviewers felt discussions were mostly useful or very useful in clarifying opinions (average is 2.14 [2.06 - 2.23] on a scale of 1 to 5). Several respondents remarked on the clarity of presentations and their importance in the evaluation:

> “It makes a big difference to hear what the reviewers that carefully examined the application thought and why when there’s some disagreement. People see different things. Bringing those things out in discussion helps assure more fairness of scores across applications when different people are reviewing.”

However, some respondents felt the distribution of comments from panel members and their relative weight on panel opinion was not even across reviewers and sometimes dependent on whether the assessment was positive or negative:

> “My experience has been that folks do not use the discussion time most effectively. The primary reviewer is accepted at face value and when a dissenting opinion is voiced, the panel seems to be reluctant to discuss and rather defers to the primary reviewer. This is a flaw that makes the review only as good as the thoroughness of ONE reviewer. And in my limited experience I have seen some shoddy reviewing. This is unfair to the team that prepares the application.”
>
> “On the whole, I felt most panel members were too polite and unwilling to offer frank opinions of weak proposals.”

As above, respondents commented on the usefulness of the opinions and interactions of unassigned reviewers in panel discussion:

> “It’s always difficult to strike a balance between having non-assigned reviewers contribute to a discussion and their relative lack of knowledge of the area. However, my opinion is that they often help to clarify points that (maybe) are obvious to the expert reviewers but probably not to the rest of the panel, resulting in a more informed final score.”

Moreover, most reviewers (79% [75%-82%]) agreed that the format and duration of the grant application discussions was sufficient to allow the non-assigned reviewers to cast well informed merit scores. However, some respondents suggested that these discussions are only effective in influencing unassigned reviewers’ scoring:

> “Discussions change very few minds among reviewers and probably are only useful to informing other, non-assigned panel members.”
>
> “While the discussion does not alter the assigned reviewers initial scores, it provides context and rationale for the scores which helps the other reviewers decide on a score.”

However, others lament that unassigned reviewers don’t read the grants, and therefore their scoring is superficially based on limited discussion, potentially leading to bias:

> “There really is not enough time for non-assigned reviewers to be able to read the grants and listen to the discussion. I think people just decide which of the assigned reviewers they like more (or whose argument they like more) and then vote with them.”
>
> “It is not clear that the content of assigned reviewer comments are driving the scoring decisions of non-assigned reviewers”
>
> “Very difficult for non-assigned panel members to actually judge the application fairly.”

Biases may be exacerbated by short discussion times, as reported by some respondents:

> “Some reviewers spend too much time presenting their review, without focusing on the most important points, leaving less opportunity for discussion.”
>
> “Discussions are usually too short, but tend to be OK.”

### Chair Facilitation

In terms of the usefulness of the chair in facilitating the application discussions, 68% of all reviewers reported that the chair’s involvement was either extremely useful or very useful (average of 2.16 [2.08-2.25] on a scale of 1 to 5). Multiple respondents remarked specifically on how their chair helped or hindered the facilitation of the discussion and nearly all who commented on the chair recognized the importance of the chair’s role in discussion and scoring:

> “The chairs of panels I have been on have been very important in directing, limiting, and policing the discussion. Most have done a poor job, even to the point of not cutting off inappropriate questions.”
>
> “A good chair is absolutely essential to promoting balanced discussion, focusing debate, not letting debate draw on when it is clear differences of opinion are not going to be resolved based on discussion. If the chair is not good at this, the study section experience can be a miserable one.”
>
> “An open-minded chair who is willing to direct discussion to key points is essential”

Several commented on the importance of the chair in facilitating participation:

> “The chair of the panel is vital for success in the review process. Having a chair that encourages discussion and differing opinions is very important to having reviewers feel their voice has been heard, their opinion is valuable and promotes continuing grant review participation”

The overwhelming majority (88% [85%-90%]) of reviewers felt the discussions were fair and balanced, and mentioned the chair’s role in ensuring balanced discussion in their comments:

> “Chair was great, fair and very well informed. It helped keep the discussion focused and helped adjust for extremes - overly laudatory reviews and extremely negative reviews. Reviewers are variable. There were a couple of reviewers who were so negative I know how the applicant must feel when reading the review. Most were balanced.”
>
> “I do know there is much effort made to provide fair and balanced discussion, though. It’s most uncomfortable when one or two members of a panel, individuals who are more vociferous or opinionated, sway quieter reviewers who actually presented more logical reasons to support their scores. In other words, scores are sometimes based on the assertiveness of the reviewers’ opinions rather than logic and rationale regarding the merits of the science. Those are situations where the Chair becomes extremely important but where I’ve seen applicants lose out.”

### Discussions and Outcome

A total of 71% of reviewers agreed that panel discussion was extremely effective or very effective in influencing the outcome of the grant (average of 2.17 [2.08-2.26] on a scale of 1 to 5). Nevertheless, many respondents suggested that there were relatively small scoring changes based on discussion:

> “The intriguing thing for me is that after a very comprehensive and high-quality discussion, in many cases the preliminary scores do not change much”

and that if outcome is affected, it is more often in a negative than positive direction:

> “Discussions rarely bring a grant to a better score, more often points out weaknesses. While I used to see that discussions brought folks to a consensus, I more often see it now as a veto of one because the only ones that achieve a fundable score are those without no negatives brought out in the discussion. This is the consequence of lots of good grants and continuing low funding lines.”

Importantly, some suggest an individual reviewer can have an undue influence on the outcome, either through dominating the discussion or through a poorly constructed initial review:

> “the discussion and influence of a reviewer varies greatly and can make or break a borderline grant. The stronger reviewer will prevail. Few are willing to get into intellectual argument, especially if they haven’t been assigned to the grant”
>
> “I have found that when the primary and secondary reviewers disagree upon initial review, discussion rarely changes the outcome much, even when one of the reviewers admits that they were wrong in down-scoring the application. And then the rest of the panel members split the difference. So, one grant with an excellent score by one reviewer and a mediocre score by the other ends up with a score outside the fundable range. And if the average of the scores places it out of the range to be discussed, the panel usually just lets it go so as to not increase the length of the meeting. So, a grant that one reviewer rated favorably can get torpedoed by a bad reviewer, even if that reviewer was totally off-base on their reasons for the bad score. Unfortunately, I have seen this from both sides - reviewer and grant applicant.”

Similarly, while 60% of reviewers definitely or most definitely agreed that the grant application discussions promoted the best science (average of 2.38 [2.30-2.46] on a scale of 1 to 5), several respondents mentioned a level of conservatism in the discussion:

> “Truly innovative grants are going to have an inherent risk. Panels often go for ‘safe’ bets.”
>
> “The dynamics of panels are always interesting. There does seem to be some level of group-think, mostly resulting in more conservative review outcomes in my experience, but it’s a complex interaction so difficult to describe accurately”

Again, respondents mentioned the disproportionate impact an individual bias can have on scoring and that unassigned reviewers may contribute significantly to bias:

> “While most reviewers are knowledgeable and unbiased, it takes just one panel member to cast doubt on a grant.”
>
> “I think the discussion with panel members who have not read the grant is pretty much a farce and can lead to dragging down of grants in an unfair manner.”
>
> “In one or two cases where others were assigned to areas that I had greater expertise in, I felt their lack of expertise led to opinionated and influential comments that swayed the panel. In this situation, when I had not read the (unassigned) grant, I was able only to comment on the correctness of the statements of the reviewer, but not offer an alternative view based on knowledge of the science under discussion. In addition, there is pressure on reviewers to make the evaluations short, which carries over to panel members not to belabor the discussion. I feel the time crunch has a negative impact on the fairness and thoroughness of the review.”

Given the above responses, we were interested to explore whether views on panel discussion and outcome were related to those of discussion facilitation. We separated respondents into 2 groups, those who felt the discussions affected the outcome (scoring 1 or 2 on this question; N=450) and those who did not feel the discussions affected the outcome (scoring 3, 4, or 5 on this question; N=184). We then compared the two groups in terms of responses surrounding discussion facilitation (**Table 1**). Significant differences were found between the two groups for all the questions, including views on reviewer participation, clarification of differing opinions, informing unassigned reviewers, and chair facilitation. In all cases, respondents who felt the outcome was affected by panel discussions viewed the discussion facilitation more favorably than those who felt the outcome was not affected by the discussions.

**Table 1.**
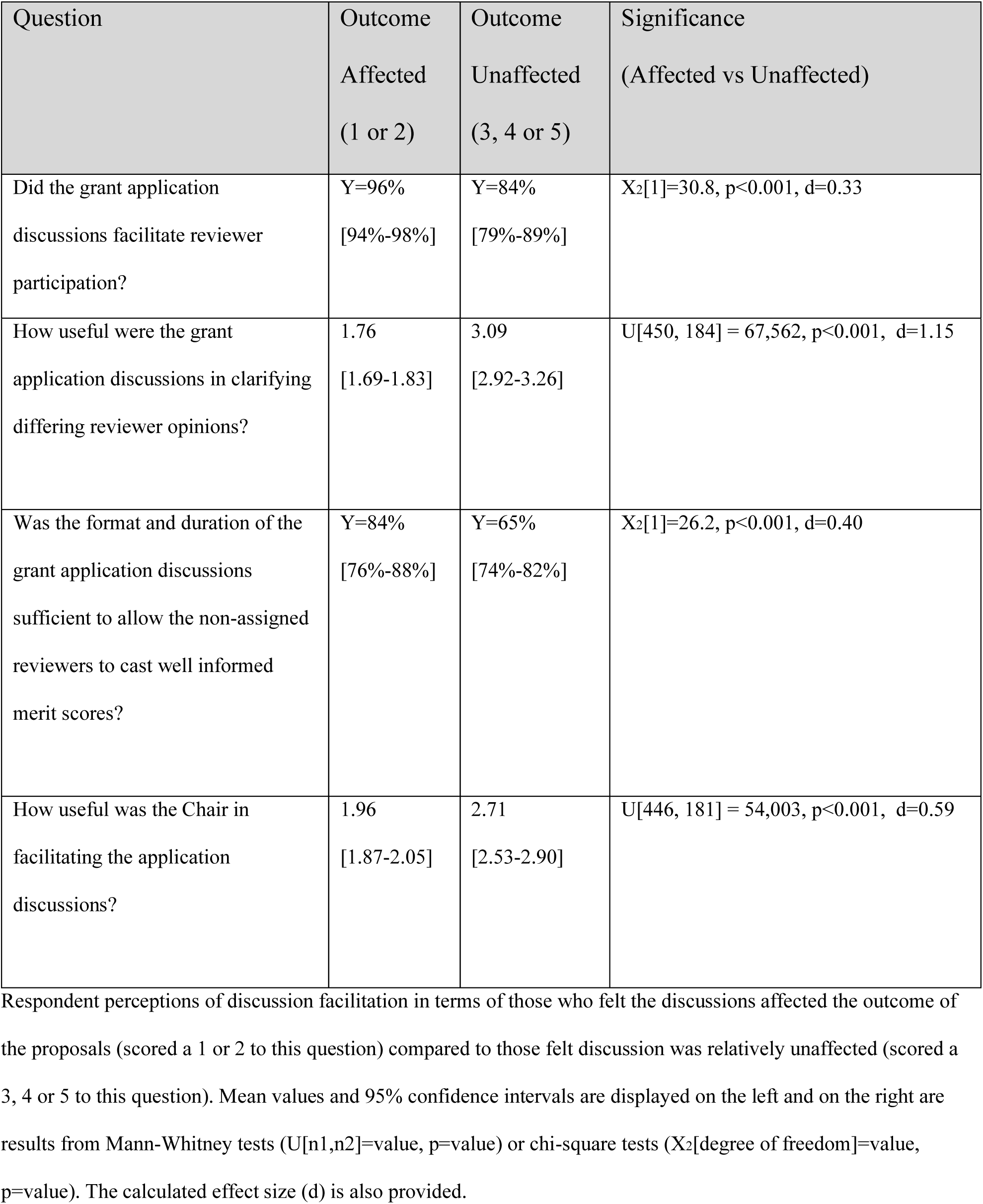
Discussion Facilitation versus Discussion Affecting Outcome.

## Discussion

Our results indicate that, in general, reviewers felt that panel discussions were well facilitated across multiple dimensions, including favorable perceptions of panel inclusivity, leadership, and quality of communication. However, our results also indicated that reviewers who do not think the discussion affects the outcome are much more likely to feel that several aspects of panel communication are problematic (**Table 1**). Respondents from this survey mentioned uneven consideration of reviewers’ opinions, low levels of participation from unassigned reviewers, and short discussion times as potential problems with panel discussions. These combined results support the idea that, at least in some instances, issues with the quality of panel discussion and panel facilitation likely limit the influence of the discussion on panel scoring.

Similarly, while the majority of respondents felt the discussions affected the outcomes, several respondents commented how such discussions contained biases that limit the fairness of the review and its ability to select the best science. In particular, some reported how an individual reviewer (if not reined in) can have a greater than intended influence on the outcome, leading to a potential source of bias during discussion. Moreover, the manner of assignments (where only a few reviewers read the application) allows for this structure to bias the outcome, particularly for unassigned reviewers’ scores. Several respondents also mentioned a level of conservatism in review panels with regards to innovative applications. While these suggestions of bias run counter to the majority of respondents who report that the discussion was fair and selected the best science, the presence of bias is still clearly an area of concern for some respondents. It may be that some reviewers are, in general, overconfident about panel effectiveness and fairness (potentially because they were directly involved in the discussion) and therefore not attuned to such biases (Moore and Healy, 2008). Future studies could gather perceptions from impartial panel observers, such as scientific review officers who manage panels for funding agencies, to determine whether such perceptions of fairness and panel effectiveness are warranted.

Importantly, most respondents recognized the importance of the chair in facilitating the panel, and many comments suggested good chairs could help elicit reviewer participation, guide balanced discussion, and play important roles in modulating the length of discussion. In addition, respondents noted the role of the chair in mitigating bias, limiting extreme reviewers, optimally leveraging panel expertise, encouraging the clear presentation of the assigned reviewer evaluations, and “directing discussion to key points.” Thus, based on the comments, there was almost universal support for the importance of the chair role in the facilitation of the panel to improve the impact of the discussion on the outcome while avoiding potential biases. Nevertheless, nearly a third of respondents did not find the chair of their most recent panel to be very effective in facilitating the application discussions. If the chair is central to the effectiveness of panel discussion, more research should focus on identifying specific facilitative behaviors of effective chairs, and specific skills that moderate discussion in inclusive and unbiased ways. Future studies of discussion quality could assess for assertive and passive personality traits and panel leadership styles (Cooke et al., 2015). For instance, variability in discussion time may be a function of chair behavior (limit-setting versus allowing discussion). Further, the effectiveness of less persuasive reviewers may be hindered by a passive chair compared to a more engaged and assertive chair. Previous research has reported the importance of score-calibration comments and even laughter in the effectiveness of panel discussion, although it is unclear if these are affected by chair facilitation (Raclaw and Ford 2017; Pier et al., 2019; Pier et al., 2017).

Our results also suggest that reviewers are unlikely to participate in discussions of proposals on which they are not the assigned reviewers, likely due to the fact that they do not read these proposals. According to some respondents, bias can result from unassigned reviewer reliance on only the discussion to inform them about proposals’ strengths and weaknesses. This model of panel scoring, where most panelists score the proposal without having read it, may not achieve the goal of leveraging panel expertise. Because many reviewers are over-burdened (Gallo 2019), solutions to achieve this goal should be explored, such as examining the optimal number of assigned reviewers (Snell 2015), or asking unassigned reviewers to read proposal abstracts and critiques ahead of the meeting.

It should be mentioned that one potential limitation to this study is the relatively low response rate (6.7%), although this rate is similar to those in other recent surveys on journal peer review (Ware 2008a, Ware 2008b, Sense about Science 2009). Furthermore, the demographics of our sample are very similar to those of NIH study section members, according to recent reports (NIH 2013). Additionally, comparing the larger, full sample of incomplete responses (n=1231) to the one used in this manuscript, we find very similar demographics, which suggests that this sample is representative of the larger population.

## Conclusions

Overall, our results find that most reviewers think the quality of panel discussion is high and does affect the outcome of the review. Conversely, our results also point to poor panel facilitation as a potential factor that limits the influence the discussion has on scoring and may even introduce biases. It is also clear from this study that reviewers feel a strong chair can help to avoid such biases and ensure engagement and inclusion; therefore, it is of great importance that future chairs should be properly trained in how to lead and facilitate a discussion. Moreover, future review processes could be informed by the science of leadership and team communication to enhance consistency, inclusivity and impartiality in panel discussions (Bos et al., 2002; Driskell et al., 2003; Kozlowski et al., 2006; Chen et al., 2013; Cooke et al., 2015).

## References

1. Bos N, Olson J, Gergle D, Olson G, & Wright Z. Effects of four computer-mediated communications channels on trust development. In Proceedings of the SIGCHI conference on human factors in computing systems. 2002; 135–140. ACM.

2. Brown AL, et al. Distributed Expertise in the Classroom. In: Salomon G, editor. Distributed Cognitions: Psychological and Educational considerations. Cambridge: Cambridge University Press; 1993. pp. 188–228.

3. Carpenter AS, Sullivan JH, Deshmukh A, Glisson SR, & Gallo SA. A retrospective analysis of the effect of discussion in teleconference and face-to-face scientific peerreview panels. BMJ open 2015; 5(9), e009138.

4. Chen G, Farh JL, Campbell-Bush EM, Wu Z, & Wu X. Teams as innovative systems: Multilevel motivational antecedents of innovation in R&D teams. Journal of Applied Psychology, 2013, 98(6), 1018.

5. Cicchetti DV. The reliability of peer review for manuscript and grant submissions: A cross-disciplinary investigation. Behavioral and brain sciences, 1991, 14(1), 119–135.

6. Cole S & Simon GA. Chance and consensus in peer review. Science. 1981; 214(4523): 881–886.

7. Cooke NJ. National Research Council. Enhancing the effectiveness of team science. National Academies Press; 2015; Jul 15

8. Driskell JE, Radtke PH, Salas E. Virtual teams: effects of technological mediation on team performance. Group Dyn. 2003; 7: 297–323.

9. Fleurence RL, Forsythe LP, Lauer M, Rotter J, Ioannidis JP, Beal A, Frank L and Selby JV. Engaging patients and stakeholders in research proposal review: the patient-centered outcomes research institute. Ann Intern Med. 2014; 161:122–30.

10. Fogelholm M, Leppinen S, Auvinen A, et al. Panel discussion does not improve reliability of peer review for medical research grant proposals. J Clin Epidemiol. 2012; 65: 47–52. doi:10.1016/j.jclinepi.2011.05.001

11. Gallo S, Thompson L, Schmaling K, and Glisson S. Risk evaluation in peer review of grant applications. Environment Systems and Decisions. 2018; 1–14.

12. Gallo SA, Carpenter AS, Glisson SR. Teleconference versus face-to-face scientific peer review of grant application: effects on review outcomes. PLoS ONE 2013; 8: e71693. doi:doi.10.1371/journal.pone.0071693

13. Gallo SA, Thompson LA, Schmaling KB, & Glisson SR. Participation and Motivations of Grant Peer Reviewers: A Comprehensive Survey. Sci Eng Ethics 2019; https://doi.org/10.1007/s11948-019-00123-1; Preprint available from: bioRxiv, 479816.

14. Kozlowski SW, & Ilgen DR. Enhancing the effectiveness of work groups and teams. Psychological science in the public interest, 2006, 7(3), 77–124.

15. Langfeldt L. The decision-making constraints and processes of grant peer review, and their effects on the review outcome. Social Studies of Science, 2001, 31(6), 820–841.

16. Liaw L, Freedman JE, Becker LB, Mehta NN, & Liscum L. Peer Review Practices for Evaluating Biomedical Research Grants. Circulation research, 2017; 121(4), e9–e19.

17. Martin MR, Kopstein A, Janice JM. An analysis of preliminary and post-discussion priority scores for grant applications peer reviewed by the Center for Scientific Review at the NIH. PLoS ONE 2010; 5:e13526. doi:10.1371/journal.pone.0013526

18. Moore DA, & Healy PJ. The trouble with overconfidence. Psychological review, 2008; 115(2): 502.

19. NIH CSR. Reviewer Quick Feedback Survey Results. 2015; https://public.csr.nih.gov/sites/default/files/2017-10/ReviewerQuickFeedbackSurveyResults.pdf (last accessed January 2019).

20. NIH OER. Enhancing Peer Review Survey Results Report. 2013;https://enhancing-peer-review.nih.gov/docs/Enhancing_Peer_Review_Report_2012.pdf (last accessed May 2019).

21. NIH. Peer Review. 2018; https://grants.nih.gov/grants/peer-review.htm (Last Accessed January 2019).

22. Obrecht M, Tibelius K, D’Aloisio G. Examining the value added by committee discussion in the review of applications for research awards. Res Eval 2007; 16: 79–91. doi:10.3152/095820207X223785

23. Pier EL, Brauer M, Filut A, Kaatz A, Raclaw J, Nathan MJ, Ford CE & Carnes M. Low agreement among reviewers evaluating the same NIH grant applications. Proceedings of the National Academy of Sciences. 2018; 115(12): 2952–2957.

24. Pier EL, Raclaw J, Carnes M, Ford CE, and Kaatz A. Laughter and the Chair: Social Pressures Influencing Scoring During Grant Peer Review Meetings. Journal of general internal medicine. 2019; Jan2:1–2.

25. Pier EL, Raclaw J, Nathan MJ, Kaatz A, Carnes M, & Ford CE. Studying the study section: How group decision making in person and via videoconferencing affects the grant peer review process. WCER Working Paper No. 2015–6. Wisconsin Center for Education Research. 2015 Oct.

26. Pier, E. L., Raclaw, J., Kaatz, A., Brauer, M., Carnes, M., Nathan, M. J., & Ford, C. E. (2017). ‘Your comments are meaner than your score’: score calibration talk influences intra-and inter-panel variability during scientific grant peer review. Research Evaluation, 26(1), 1–14.

27. Raclaw J and Ford CE. Laughter and the management of divergent positions in peer review interactions. Journal of pragmatics. 2017; 113: 1–15.

28. Sense About Science “Peer Review Survey” 2009 http://archive.senseaboutscience.org/pages/peer-review-survey-2009.html (last accessed May 2019)

29. Snell RR. Menage a quoi? Optimal number of peer reviewers. PloS one 2015, 10(4), e0120838.

30. Vo NM and Trocki R. Virtual and Peer Reviews of Grant Applications at the Agency for Healthcare Research and Quality. South Med J. 2015; 108(10): 622–6.

31. Ware M, & Monkman M. Peer review in scholarly journals: Perspective of the scholarly community—An international study. London, UK: Publishing Research Consortium. 2008a

32. Ware Mark. Peer review: benefits, perceptions and alternatives. Publishing Research Consortium 2008b: 4.

